# Primary saturation of α, β-unsaturated carbonyl containing fatty acids does not abolish electrophilicity

**DOI:** 10.1101/2020.09.24.311985

**Authors:** Nathaniel W. Snyder, James O’Brien, Bhupinder Singh, Gregory Buchan, Alejandro D. Arroyo, Xiaojing Liu, Anna Bostwick, Erika L. Varner, Anusha Angajala, Robert W. Sobol, Ian A. Blair, Clementina Mesaros, Stacy G. Wendell

## Abstract

Metabolism of polyunsaturated fatty acids results in the formation of hydroxylated fatty acids that can be further oxidized by dehydrogenases, often resulting in the formation of electrophilic, α,β-unsaturated ketone containing fatty acids. As electrophiles are associated with redox signaling, we sought to investigate the metabolism of the oxo-fatty acid products in relation to their double bond architecture. Using an untargeted liquid chromatography mass spectrometry approach, we identified mono- and di-saturated products of the arachidonic acid-derived 11-oxoeicosatetraenoic acid (11-oxoETE) and mono-saturated metabolites of 15-oxoETE and docosahexaenoic acid-derived 17-oxodocosahexaenoinc acid (17-oxoDHA) in both human A549 lung carcinoma and umbilical vein endothelial cells. Notably, mono-saturated oxo-fatty acids maintained their electrophilicity as determined by nucleophilic conjugation to glutathione while a second saturation of 11-oxoETE resulted in a loss of electrophilicity. These results would suggest that prostaglandin reductase (PTGR1), known for its reduction of the α,β-unsaturated double bond, was not responsible for the saturation of oxo-fatty acids. Surprisingly, knockdown of PTGR1 expression by shRNA confirmed its participation in the formation of 15-oxoETE and 17-oxoDHA mono-saturated metabolites. Furthermore, overexpression of PTGR1 in A549 cells increased the rate and total amount of oxo-fatty acid saturation. These findings will further facilitate the study of electrophilic fatty acid metabolism and signaling in the context of inflammatory diseases and cancer where they have been shown to have anti-inflammatory and anti-proliferative signaling properties.

**Highlights:**

- Primary saturation of electrophilic fatty acids does not abolish biological activity.
- Prostaglandin reductase 1 reduces double bonds in fatty acids that are structurally similar to 15-keto-prostaglandin E_2_.
- Prostaglandin reductase 1 reduces non-carbonyl adjacent double bonds.

## 1. Introduction

Omega-6 (Ω-6) and omega-3 (Ω-3) polyunsaturated fatty acids (PUFA) are enzymatically metabolized by cyclooxygenases (COX), lipoxygenases (LO), and cytochrome P450s or oxidized by reactive oxygen and nitrogen species to numerous fatty acid metabolites, many of which are biologically active (1–4). Metabolism produces a diversity of structural alterations, some subtle, which result in a plethora of signaling actions. Initial hydrogen abstraction and oxygenation of parent fatty acids, such as arachidonic acid (AA, Ω-6) and docosahexaenoic acid (DHA, Ω-3) forms hydroperoxy fatty acids that are quickly reduced by peroxidases to their corresponding hydroxy metabolites, the AA-derived hydroxyeicosatetraenoic acids (HETEs) and the DHA-derived hydroxydocosahexaenoic acids (HDoHEs) (3, 5, 6). One predominant pathway of hydroxy fatty acid metabolism is oxidation by dehydrogenases, such as 15-hydroxyprostaglandin dehydrogenase (15PGDH), to their corresponding ketone-containing oxo-fatty acid products (7–9). These products contain an α,β-unsaturated carbonyl that presents electrophilic properties. Unlike many bioactive fatty acids, electrophilic oxo-fatty acid signaling, with the exception of 5-oxoeicosatetraenoic acid (oxoETE), is G-protein coupled receptors (GPCR) independent. Rather, electrophilic fatty acids form covalent Michael addition adducts with nucleophilic amino acids, such as cysteine, that are found in critical redox transcriptional regulatory proteins and pro-inflammatory enzymes (6, 10–12). Electrophilic α,β-unsaturated ketone containing fatty acids derived from Ω−6 and Ω-3 PUFA have anti-proliferative, anti-inflammatory, and redox signaling properties (13–16). While a great deal of focus has been placed on the formation of electrophilic protein adduction and signaling (17, 18); very little is known about the downstream metabolism of these electrophilic fatty acids.

Based on the exemplary metabolism of 15-ketoprostaglandin E_2_ (15-ketoPGE_2_) to 13, 14-dihydro-15-ketoPGE_2_, it is reasonable to hypothesize that other α,β-unsaturated carbonyl-containing electrophiles would follow a similar pathway of metabolism. 15-ketoPGE_2_ is formed from the conversion of AA to PGH_2_ by COX followed by prostaglandin E synthase formation of PGE_2_. PGE_2_ contains a hydroxyl group at C15 that is oxidized by 15PGDH to form 15-ketoPGE_2_, an electrophile with bioactive signaling capabilities (19). Inactivation of the electrophilic moiety occurs via reduction of the α,β-unsaturated bond by prostaglandin reductase (PTGR)1, commonly referred to as leukotriene B_4_ 12-hydroxydehydrogenase, 2-alkenal reductase, or alkenal/one oxidoreductase in humans and dithiolethione inducible gene in rat (7). Electrophile formation and subsequent reduction by PTGR1 is documented for the AA-derived Lipoxin A_4_ as well other prostaglandins and alkenals including 4-hydroxynonenal (20, 21). PTGR1 also reduces the double bond of the electrophilic nitroalkene, nitro-oleic acid, resulting in the formation of the non-electrophilic nitro-alkane, nitro-stearic acid (22).

Based on these data, one would assume that other α,β-unsaturated carbonyl-containing electrophiles, such as the oxoETEs and the oxodocosahexaenoic acids (oxoDHAs) would also be inactivated by PTGR1. In this current study, the metabolic fate of two oxoETEs derived from arachidonic acid, 11- and 15-oxoETE and the DHA-derived 17-oxoDHA, was investigated. We implicate PTGR1 (although not exclusively responsible for) in the saturation of the oxoFAs 11-, 15-oxoETE and 17-oxoDHA by purified enzyme activity, as well as knockdown and overexpression experiments. Interestingly and contrary to our original hypothesis, we show that the mono-saturated metabolite of all three oxo-fatty acids retains electrophilicity.

## 2. Materials and Methods

### 2.1 Reagents and materials

11-oxoETE, 15-oxoETE, and the [^13^C_20_]-15-oxoETE stable isotope internal standard (ISTD) were synthesized in-house as previously reported (11). Peroxide free arachidonic acid, 17-oxoDHA, 5-oxoETE-d_7_, 15-ketoPGE_2_, 13,14-dihydro-15-ketoPGE_2_, and 13,14-dihydro-15-ketoPGE_2_-d_4_ were purchased from Cayman Chemical (Ann Arbor, MI). Ammonium formate, glacial acetic acid, β-mercaptoethanol and dimethyl sulfoxide (DMSO) were purchased from Sigma-Aldrich (St. Louis, MO). HPLC grade chloroform as well as Optima LC/MS grade methanol, acetonitrile, water, isopropanol, ammonium acetate, and formic acid were purchased from Fisher Scientific (Pittsburgh, PA). Ham’s F-12K media, Medium 200, HBSS, low serum growth supplement (LSGS) kit, human umbilical vein endothelial cells (HUVECs), streptomycin, and penicillin were purchased from Invitrogen (Carlsbad, CA). Fetal bovine serum (FBS) was from Gemini Bioproducts (West Sacramento, CA). Human lung adenocarcinoma A549 cells (CRM-CCL-185) were obtained from ATCC (Manassas, VA).

### 2.2 Cell treatment with oxo-fatty acids

HUVECs were grown until 80% confluence on Collagen IV coated plates from Becton Dickinson (Bedford, MA), and maintained in medium 200 supplemented with LSGS. Cells were treated with 10 μM 11-oxoETE, 15-oxoETE in 0.25% DMSO containing media. A549 cells were grown to 80% confluency in Ham’s F12K media containing 10% FBS and 1% pen/strep. All cells were maintained at 37 °C with 5% CO_2_. At the time of treatment, media was replaced with HBSS and 10 μM 15-oxoETE or 17-oxoDHA in 0.25% DMSO was added to A549 cells. An initial treatment time of 90 minutes at 37°C was chosen based on previously published uptake data (14, 23). In targeted experiments, either 2.5 ng of [^13^C_20_]-15-oxoETE or 20 ng of 5-oxoETE-d_7_ as indicated was added as an internal standard (ISTD) and the extraction was performed as previously described with minor modifications (24). Briefly, cells were lifted with a scraper, mixed, and an aliquot of the resulting suspension was transferred to a glass tube. Folch extractions were performed with 2 volumes of 2:1 CHCl_3_:MeOH with 0.1% acetic acid. The organic phase was evaporated to dryness under N_2_. The samples were dissolved in 100 μL 70:30 H_2_O:CH_3_CN with 0.1% formic acid and filtered through Costar centrifuge filters (0.22 μm nylon) before transfer into HPLC vials for LC-HRMS analysis.

### 2.3 HUVEC Reduction Time Course

Media containing 0.25% DMSO with 10 μM 11-oxo-ETE, 11-oxoETE-ME, 15-oxoETE, or 15-oxoETE-ME was added to HUVECs and the cells were incubated at 37°C for 5 min, 30 min, 90 min, 3 hr, 6 hr or 9 hr. At each time point, 450 μL of the supernatant media was collected and transferred to a glass tube. The cells were gently scraped in the remaining 450 μL of media and transferred to a glass tube. 2.5 ng of [^13^C_20_]-15-oxoETE ISTD was added followed by subsequent Folch extraction as previously described (24). The organic phase was evaporated to dryness under N_2_. The samples were dissolved in 100 μL of 70:30 H_2_O:CH_3_CN with 0.1% formic acid then filtered using 0.22 μm nylon centrifuge filters and transferred into HPLC vials for LC-high resolution mass spectrometry (LC-HRMS) analysis.

### 2.4 A549 Reduction Time Course

A549 WT cells, A549 Scramble, A549 PTGR1 KD and PTGR1 OE cells were treated with 10 uM 15-oxoETE, 10 uM 17-oxoDHA or 10 uM 15-ketoPGE_2_ (positive control) in 12 well plates when cells were at 80% confluency. The treatments lasted for 6 h with 5 collection time points (5 min, 30 min, 90 min, 3 hr, and 6 hr). Cell lysates were scraped in 1 mL of PBS and the internal standard, 5-oxoETE-d_7_ (20 ng) was added before samples were extracted with 5 mL of CHCl_3_:MeOH (2:1). Samples were vortexed and centrifuged at 2800 x *g* for 10 min. The organic layer was transferred to a new tube, dried under N_2_ and stored at −80°C before analysis.

### 2.5 PTGR1 shRNA knockdown

Lentiviral particles were generated by co-transfection of 4 plasmids [Control plasmid (pLKO.1-SCRshRNA-Puro) or one of the five different PTGR1-specific shRNA expressing plasmids, pLKO.1-shRNA-PTGR1.1-5 together with pMD2.g (VSVG), pVSV-REV and pMDLg/pRRE] into 293-FT cells using TransIT®-2020 Transfection reagent, with support from the UPCI Lentiviral Facility. The PTGR1-specific shRNAs target the following sequences in the PTGR1 gene: (1) CGTCTCCTGATGGTTATGATT, (2) GCCTACTTTGGCCTACTTGAA, (3) CTATCCTACTAATAGTGACTT, (4) CTTGGATTTGATGTCGTCTTT, and (5) GACTTGCTGAAATGGGTCTTA (Table 1). The collection and isolation of lentiviral particles and A549 cell transduction were performed as described previously (9). Stable cell lines were selected in puromycin (1 μg/ml) for 2 weeks. Knockdown (KD) of PTGR1 was confirmed by immunoblot and checked throughout experimentation. For PTGR1-KD cells, proteins were normalized to 15 μg/μL and each sample was loaded onto NuPAGE 4-12% Bis-Tris Midi Gel. Proteins on the gel were then transferred to nitrocellulose membrane and blocked in 5% non-fat dried milk in TBST for 1 h. The immunoblot was probed with 1:1000 PTGR1 rabbit antibody for 24 h (Aviva Systems Biology, San Diego, CA) and a horseradish peroxidase-conjugated anti-rabbit secondary antibody for 1 h. Membranes were visualized using BioRad Chemi-doc (Hercules, CA)

**Table 1:**
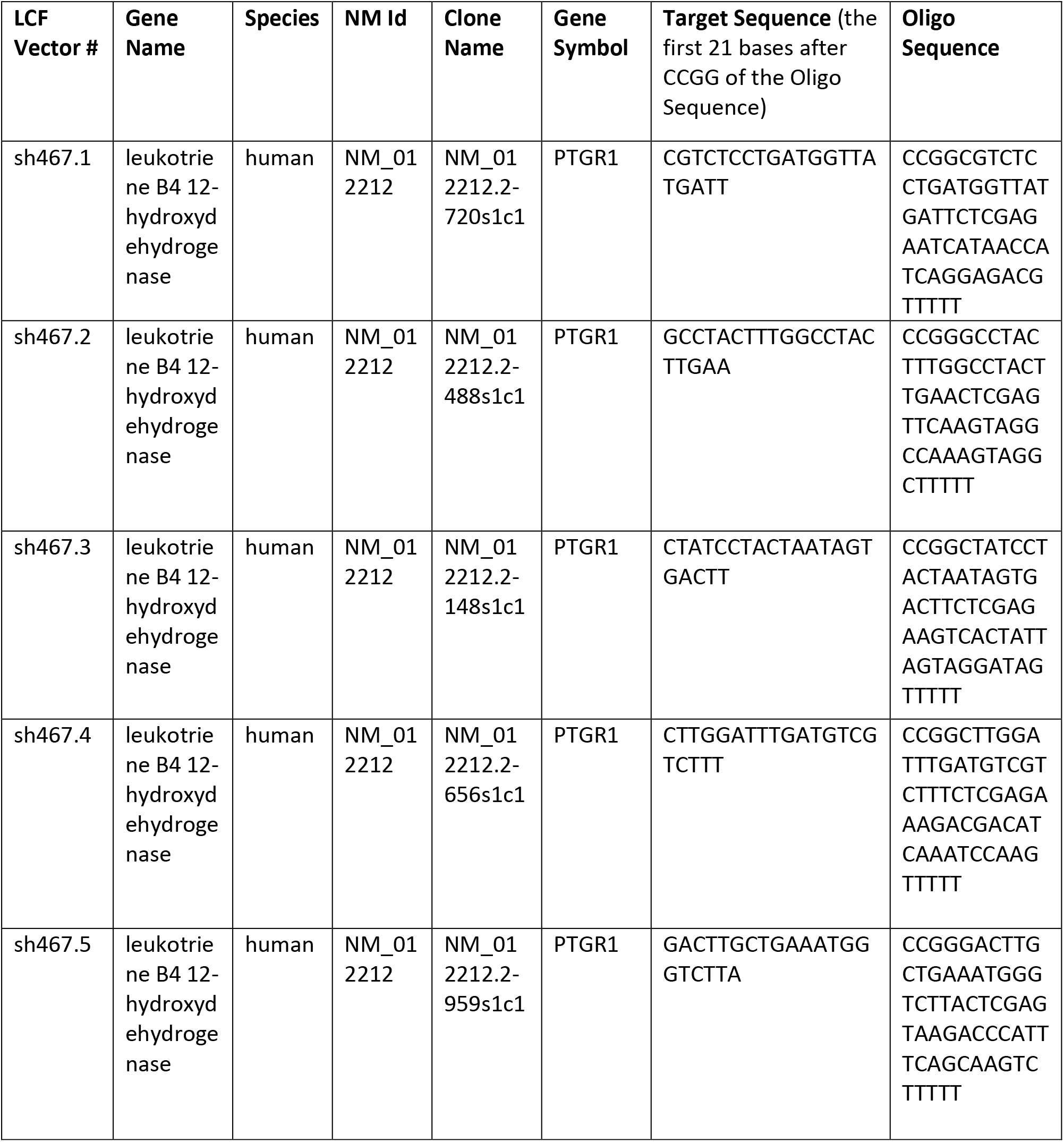
PTGRI shRNA sequences.

### 2.6 Vector development for over-expression of PTGR1

Lentiviral vectors were developed for expression of human prostaglandin reductase 1 (hPTGR1) with an N-terminal 3xFLAG tag (lab stock#2040; pLV-Puro-EF1A-3xFLAG-hPTGR1) and a control vector with a non-coding ‘stuffer’ fragment replacing the 3xFLAG-hPTGR1 open reading frame (lab stock#2039; pLV-Puro-EF1A-Stuffer), each prepared by Vector Builder Inc.

### 2.7 Lentivirus production and cell transduction for over-expression of PTGR1 in A549 cells

293FT cells were cultured using the following media: DMEM high glucose (Corning cat#15-017-CV) supplemented with 10%FBS, 0.1mM MEM non-essential amino acids NEAA (Gibco cat#1140-050); 6mM L-Glutamine (Gibco cat# 25030-081); 1mM Sodium Pyruvate (Gibco cat#11360-070) and 1%Pen-Strep (Gibco cat#15140-122). Lentiviral particles were generated by co-transfection of 4 plasmids into 293FT cells using TransIT-X2 Transfection reagent: the packaging vectors pMD2.g(VSVG), pRSV-REV, and pMDLg/pRRE together with the appropriate shuttle vectors (pLV-Puro-EF1A-3xFLAG-hPTGR1 or pLV-Puro-EF1A-Stuffer). Forty-eight hours after transfection, lentivirus-containing supernatant was collected and passed through 0.45 mM filters to isolate the viral particles as described previously (25, 26)(25, 26). A549 cells were cultured using the following media: DMEM high glucose (Corning cat#15-017-CV) supplemented with 10%FBS; 2mM L-Glutamine (Gibco cat#25030-081) and 1%Pen-Strep (Gibco cat#15140-122). Lentiviral transduction was performed as follows: A549 cells (1-2×10^5^) were seeded into 6-well plates. Twenty-four hours later, lentiviral particles were mixed with polybrene (2μg/ml) and then added to the cells (1ml). Cells were incubated at 32°C overnight and then the medium with lentiviral particles was removed and replaced with fresh medium. The transduced A549 cells were then cultured for 48 hr at 37°C before selection with puromycin (1μg/ml) for 2 weeks. Four separate transductions were carried out for each vector, giving rise to four control cell lines (A549/2039.1−.4) and four cell lines over-expressing PTGR1 (A549/3xFLAG-PTGR1.1−.4).

#### Immunoblot for validation of PTGR1 over-expression

Cell protein extracts (whole cell lysates, WCL) were prepared from A549/2039 cells (empty vector controls) and A549/3xFLAG-PTGR1 cells as follows. Briefly, cells were seeded into a 60-mm cell culture dish. After reaching 75-80% confluency, cells were washed twice with cold PBS, collected, and lysed with an appropriate volume of 2x clear Laemmli buffer (2% SDS, 20% glycerol, 62.5mmol/l Tris-HCl pH6.8). Cell lysates were boiled for 10 min and quantified with the DC protein assay kit following the microplate protocol provided by the company (Bio-Rad). Whole cell lysates (15-40μg protein) were loaded onto precast NuPAGE® Novex® 4-12% Bis-Tris gels, run 1hr at 120V. Gel electrophoresis separated proteins were transferred onto a PVDF membrane using a Turboblotter (Bio-Rad). The membrane was first blocked with B-TBST (TBS buffer with 0.05% Tween-20 and supplemented with 5% blotting grade non-fat dry milk; Bio-Rad) for 1 hr at room temperature and subsequently blotted with the primary antibodies in B-TBST overnight at 4°C. The primary antibodies and dilutions used are as follows: anti-PTGR1 (Aviva Systems Biology, cat#50-014-61350; 1:1000); anti-FLAG M2 (Sigma, cat#F1804; 1:1000); loading control anti-⍺-Tubulin (Invitrogen, cat#62204; 1:1000). After washing, membranes were incubated with secondary antibodies in B-TBST for 1 hr (room temperature). The following HRP conjugated secondary antibodies were used: Bio-Rad Goat anti-mouse HRP conjugate (Cat#1706516; 1:1000) and Bio-Rad anti-rabbit HRP conjugate (Cat#1706515; 1:1000). After washing, the membrane was illuminated with a chemiluminescent substrate. Protein bands were imaged using a Bio-Rad Chemi-Doc MP imaging system.

### 2.8 LC-MS discovery and validation of oxo-fatty acid metabolites

For LC-HRMS analysis, LC separations were conducted as previously described (22) using a Waters nano-ACQUITY UPLC system (Waters Corp., Milford, MA, USA). A Waters XBridge BEH130 C18 column (100 μm × 150 mm, 1.7 μm pore size; Waters Corp) was employed for reversed phase separation with a 3 μL injection. For gradient 1, used during discovery and targeted validation experiments, the flow rate was 1.8 μL/min, solvent A was 98:2 water/acetonitrile (v/v) with 0.1% formic acid, and solvent B was 99.5:0.5 acetonitrile/water (v/v) with 0.1% formic acid. The gradient was as follows: 2% B at 0 min, 2% B at 3 min, 3.6% B at 11 min, 8% B at 15 min, 8% B at 27 min, 50% B at 30 min, 50% at 35 min, and 2% B at 37 min. Separations were performed at 30°C. For gradient 2, used for the confirmation of MS/HRMS and adduct studies with GSH, the flow rate was 1.5 μL/min, solvent A was 4:6 water/acetonitrile (v/v) with 0.1% formic acid and 10 mM ammonium formate and solvent B was 1:9 acetonitrile/isopropanol (v/v) with 0.1% formic acid and 10 mM ammonium formate. The gradient was as follows: 30% B at 0 min, 30% B at 6 min, 60% B at 11 min, 60% B at 15 min, 80% B at 27 min, 80% B at 30 min, 85 % at 35 min, 85 % at 45 min, 30 % at 47 min, and 30% B at 55 min. Separations were performed at 50°C. HRMS analysis was conducted using an LTQ XL-Orbitrap hybrid mass spectrometer (Thermo Fisher) in either positive or negative ion mode with a Michrom captive spray ESI source as previously described (22). For gradient 1 from above, the operating conditions were: spray voltage at 4 kV; capillary temperature at 250°C; capillary voltage at 35 V, tube lens 60 V. For gradient 2, the operating conditions were: spray voltage at 1.5 kV; capillary temperature at 200°C; capillary voltage at 0 V, tube lens 80 V. Analysis of differential abundant features was carried out by SIEVE 2.0, Metaboanalyst 2.0, and XCMS (26,27). Metabolic discovery experiments conducted in HUVEC cells revealed a diversity of differentially abundant features from the lipid fraction as demonstrated by the respective cloud plots generated with XCMS.

#### LC-MS/MS Characterization in A549 Cells

For the analysis of 15-oxoETE and 17-oxoDHA metabolism in A549 cells with and without PTGR1 shRNA KD and Scr shRNA control the samples were run in negative ion mode on a Sciex 6500 triple quadrupole ion trap coupled to a Shimadzu LC controller/pump and CTC Pal autosampler. A Phenomenex Luna C18(2), 2 × 100 mm was used for separation with a gradient of H_2_O + 0.1% acetic acid (A) and ACN + 0.1% acetic acid (B) using a flow rate of 0.6 mL/min. Solvent B started at 30% and increased to 100% B over 11 min. B was held at 100% for 3 min before returning to initial conditions to equilibrate the column for 2 min. The α, β-unsaturated ketones and metabolites were analyzed using the following source conditions: DP −65, EP −10, CE-25, and CXP −12. The MS source parameters were the following: CAD 11, curtain gas 30, gs1 80, gs2 85, ISV-4500 and source temp 600 °C. The following transitions were monitored: 15-oxoETE: 317.2→113.2, dihydro-15-oxoETE: 319.2→221.2, 15-HETE: 319.2→219.2, 5-oxoETE-d_7_: 324.2→210.2, 17-oxoDHA: 341.2→297.2, 17-HDoHE: 343.2→273.2. dihydro-17oxoDHA: 343.2→299.2, 15-ketoPGE_2_: 349.2→161.2, 13,14-dihydro-15-ketoPGE_2_: 351.2→235.2, and 13,14-dihydro-15-ketoPGE_2_-d_4_: 355.2→ 239.2. Confirmation of the position of double bond saturation was performed on a Thermo Fisher Q Exactive coupled to a Vanquish UHPLC using the same gradient described above. Samples were analyzed using full scan and parallel reaction monitoring, at a resolution of 35K and fragment ions were confirmed using accurate mass < 5ppm. For quantification of parent oxoFAs and metabolites in the recombinant protein, knockdown and overexpression experiments the same LC method was used on a Ultimate 3000 UHPLC coupled to a Q Exactive Plus.

### 2.9 Statistics

For the untargeted metabolomics analysis, a ranked list by fold change versus control using a *t*-test (Welch’s unpaired) and a p-value cut-off of p < 0.01 was generated. Time courses are plotted using SEM and multiple paired t-test are used to determine significance in PTGR1 KO and OE experiments. The p-value and replicate number are listed within the figure legends.

## 3. Results

### 3.1 Untargeted discovery reveals distinctive patterns of oxoETEs metabolism in HUVECs

To determine possible routes of 11-oxoETE and 15-oxoETE metabolism, HUVECs were treated for 90 min with 10 μM of each compound. LC-HRMS was conducted in an untargeted manner on the extracted organic (lipid) and aqueous fractions. Analysis of differential abundant features was carried out and the top 25 features based on fold change and p-value from each analysis were manually curated and representative chromatograms were extracted from raw data samples (www.xcmsonline, Job #1198294 and 1198300). Two sequential mass shifts of +2 amu (proposed mono- and di-saturated species) and an increased retention time were chosen for follow up targeted analysis for 11-oxoETE (Fig. 1A), as this pathway of metabolism has not been reported previously. Only one +2 amu shift (mono-saturated) was observed for 15-oxoETE (Fig. 1B). These putative 15-oxoETE metabolites were confirmed by comparison to the treatment of HUVECs with [^13^C_20_]-15oxoETE (Fig. 1C). This pattern of retention time and mass shift seems logically indicative of a reduction in double bonds (Scheme 1). Accurate mass matched predicted values for both parent and saturated compounds for 11-oxoETE and 15-oxoETE in positive and negative ion mode within 2.5ppm, so a 5ppm window was selected to extract all ion chromatograms.

**Figure 1:**
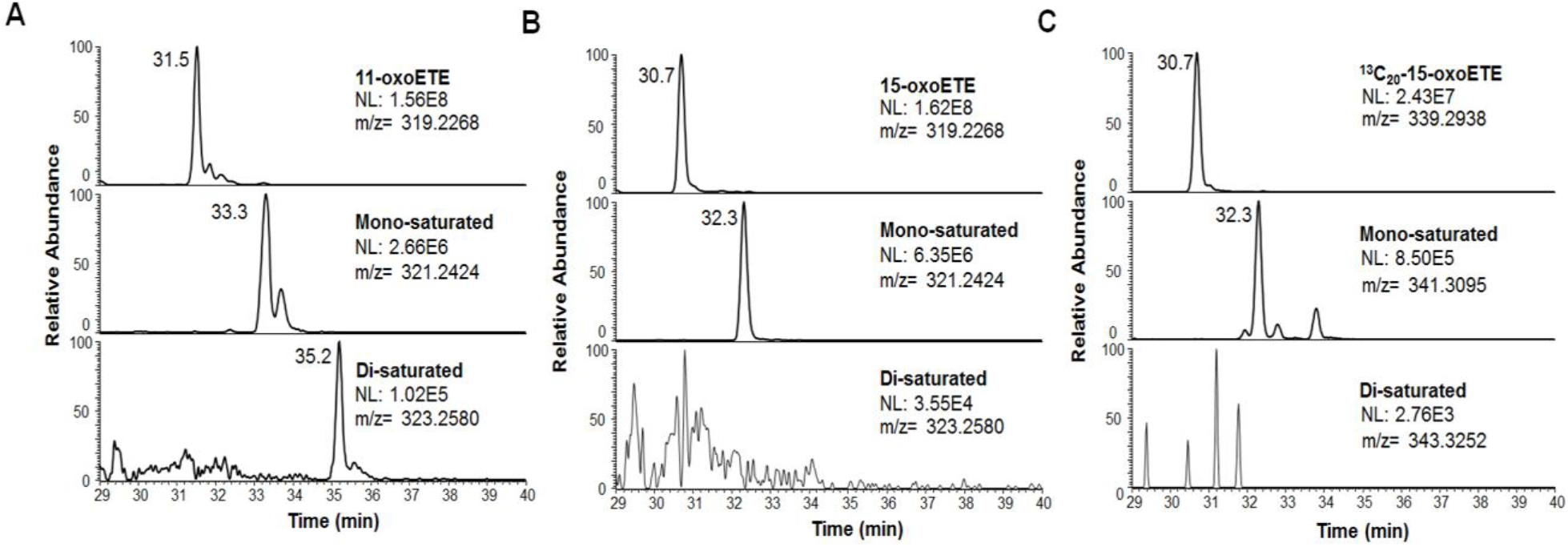
Extracted ion chromatograms (XIC) of LC-HRMS analysis of lipid extract from oxoETE treated HUVECs. Mass spectral features from the discovery experiment were extracted from representative chromatograms with a 5 ppm mass window from the given *m/z* from positive ion mode LC-HRMS analysis of lipid extracts of HUVECs treated with (A) 11-oxoETE, (B) 15-oxoETE, (C) [^13^C_20_]-15-oxoETE. Peaks putatively corresponding to double bond saturation were apparent for one and two saturations in 11-oxoETE. Only one saturation was detectable for 15-oxoETE and [^13^C_20_]-15-oxoETE.

### 3.2 Time course studies reveal accumulation of saturated metabolites

HUVECs were treated as described above, using 11-oxoETE or 15-oxoETE over 9 hr. The area of the target peak was integrated and expressed as a ratio over the area of the [^13^C_20_]-15-oxoETE ISTD peak that was spiked into the samples immediately before extraction. Values were expressed as the ratio of analyte area/ISTD for each time point. For 11-oxoETE, both mono-(Fig. 2A), and di-saturated (Fig. 2B) products appeared at the 5 min time point and accumulated over time in cell lysate (Fig. 2). Accumulation over time of the mono-saturated 15-oxoETE (Fig. 2C) metabolite was similarly observed.

**Figure 2:**
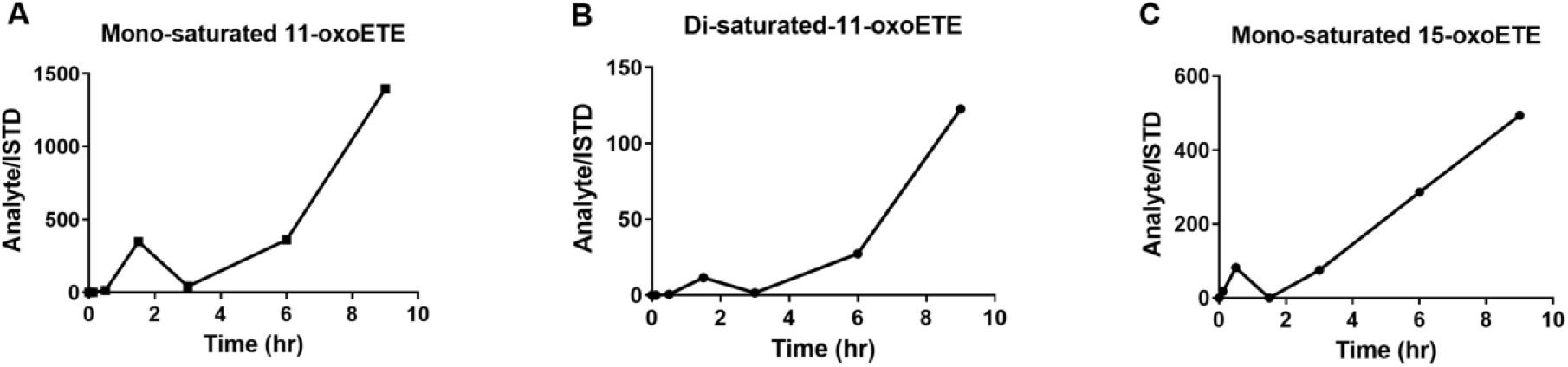
Time course of oxoETE metabolite accumulation in HUVECs. Accumulation of putative metabolites after treatment with 10 μM 11-oxoETE or 15-oxoETE. Area under the curve was calculated from the XIC of metabolites. [^13^C_20_]-15-oxoETE was used as the internal standard and to normalize for extraction and ionization efficiency for relative quantitation. Values are expressed in area under the peak for (Analyte/ISTD) for the (A) mono-saturated 11-oxoETE metabolite, the (B) di-saturated 11-oxoETE metabolite and the (C) mono-saturated 15-oxoETE metabolite.

### 3.3 MS/MS of parent oxoETEs and saturated metabolites

LC-MS/MS with collision induced dissociation (CID) fragmentation was utilized to elucidate the structure of the 11-oxoETE (Fig. 3A) and its corresponding metabolites. Under the described LC-MS conditions, 11-oxoETE (*m/z* 317.2122) eluted at 5.5 minutes and extracted MS2 fragment ion species of *m/z* 299.2015, *m/z* 273.2222, and *m/z* 211.1341 also demonstrated corresponding peaks at this retention time (Fig. 3B). As predicted, mono-saturated 11-oxoETE (*m/z* 319.2273) (Fig. 3C) and di-saturated 11-oxoETE (*m/z* 321.2429) (Fig. 3D) species eluted at later retention times on the reversed phase column. Corresponding diagnostic MS2 fragment ions were +2 (*m/z* 301.2172 and *m/z* 275.2380) and +4 amu (*m/z* 303.2328 and *m/z* 277.2535) higher than those for 11-oxoETE. The 11-oxoETE MS/MS spectrum exhibited *m/z* 165.1286 (δ = 0.6ppm) that is formed by α-cleavage to the carbonyl at C11 (Fig. 3B). The MS/MS spectrum of the mono-saturated 11-oxoETE retained the fragment at *m/z* 211.1341, but the *m/z* 165.1286 was absent, instead *m/z* 167.1443 was present, indicating that the first saturation was likely at the C14-C15 double bond (Fig. 3C). The *m/z* 211.1341 was not present in the di-saturated metabolite and the *m/z* 167.1443 was still present (Fig. 3D) thus inferring that the electrophilic double bond was saturated. However, further experiments were needed to differentially determine a saturation at C12-C13 vs C14-C15 for the mono- and di-saturated metabolites, with critical implications for the electrophilic character of 11-oxoETE.

**Figure 3:**
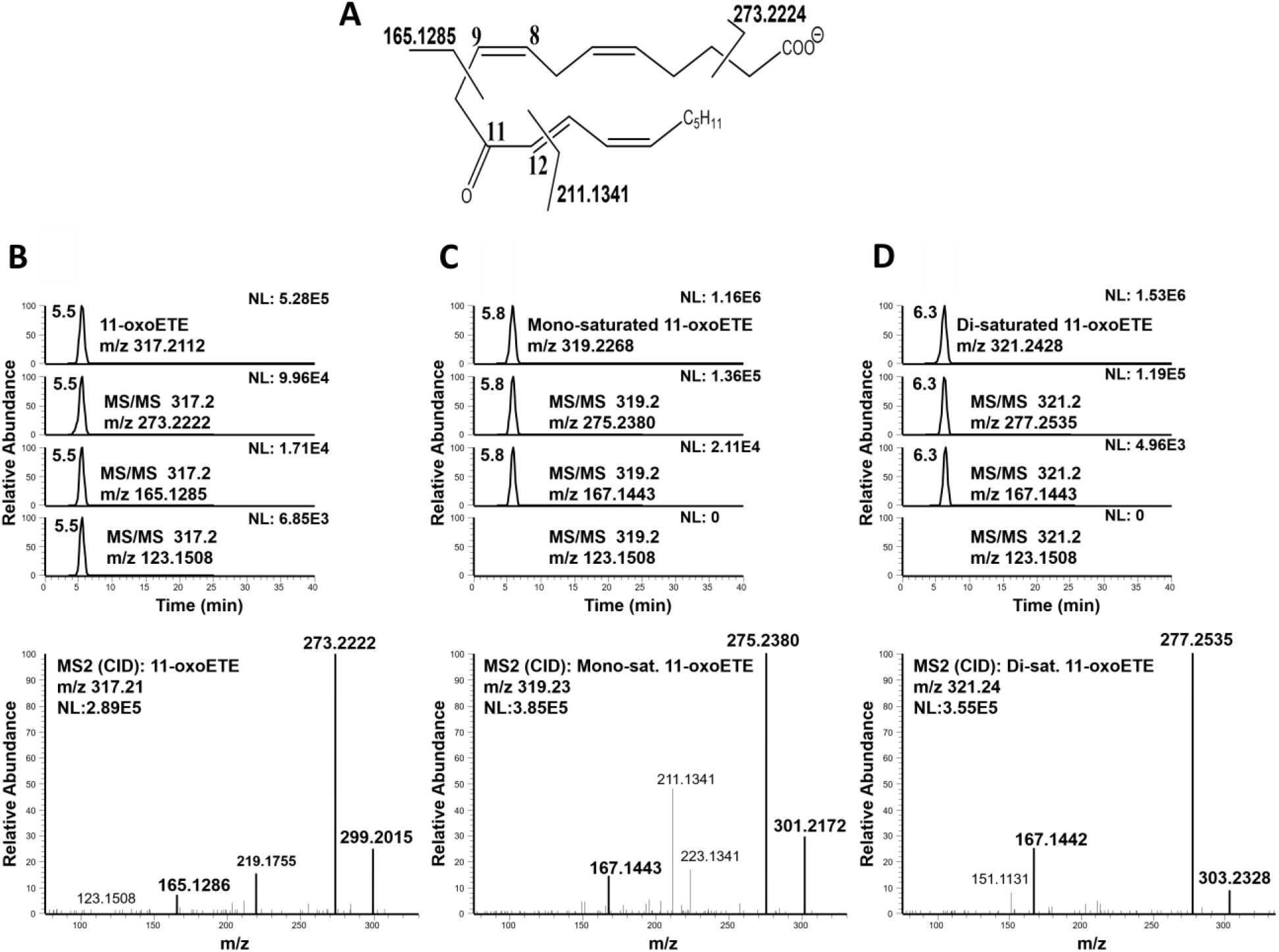
LC-MS/HRMS confirmation of 11-oxoETE saturation products. (A) 11-oxoETE structure and diagnostic fragmentation. Extracted chromatograms and product ion spectrum in negative ion mode for (B) 11-oxoETE (C) the mono-saturated 11-oxoETE metabolite, and (D) the di-saturated 11-oxoETE metabolite.

The negative ion MS/MS spectrum of 15-oxoETE (Fig. 4A) was identical with published CID spectrum by MacMillan, *et. al* (28) and diagnostic fragment ions at *m/z* 219.1755, *m/z* 203.1080, and *m/z* 113.0977 (Fig. 4B) at a retention time of 5.1 min. As would be expected from the published spectra, the MS/MS spectrum of the mono-saturated 15-oxoETE contained fragment ions with a mass increase of +2 amu compared to 15-oxoETE, including *m/z* 319.2268, *m/z* 221.1545, and *m/z* 205.1236 at a later retention time of 5.4 min (Fig. 4C). The best described fragmentation of 15-oxoETE is due to the breaking of the double bond allylic to the carbonyl moiety. This generates the fragments *m/z* 113.0977, corresponding to the part of the molecule without any other double bonds, and *m/z* 219.1394, corresponding to the part of the molecule containing all of the remaining double bonds (Fig. 4A). The MS/MS spectrum of the mono-saturated 15-oxoETE did show an intense fragment at 113.0977 strongly suggesting that the first reduction did not occur at C13-C14. The fragment corresponding to the *m/z* 219.1394 shifted to *m/z* 221.1549 is consistent with the first reduction taking place at either the C5-C6, C8-C9 or C11-C12 double bond. However, we can postulate that saturation of 15-oxoETE to form the mono-saturated product likely occurs at the C5-C6 double bond as the fragment at *m/z* 165.1273, corresponding to the chemical composition C_11_H_17_O found in 15-oxoETE is also accompanied by a fragment at *m/z* 167.1066 in the mono-saturated product (Fig. 4D). This fragment corresponds to the chemical composition C_10_H_15_O_2_, a fragment adjacent to the C8-C9 double bond that would likely not be present if saturation occurred at this position.

**Figure 4:**
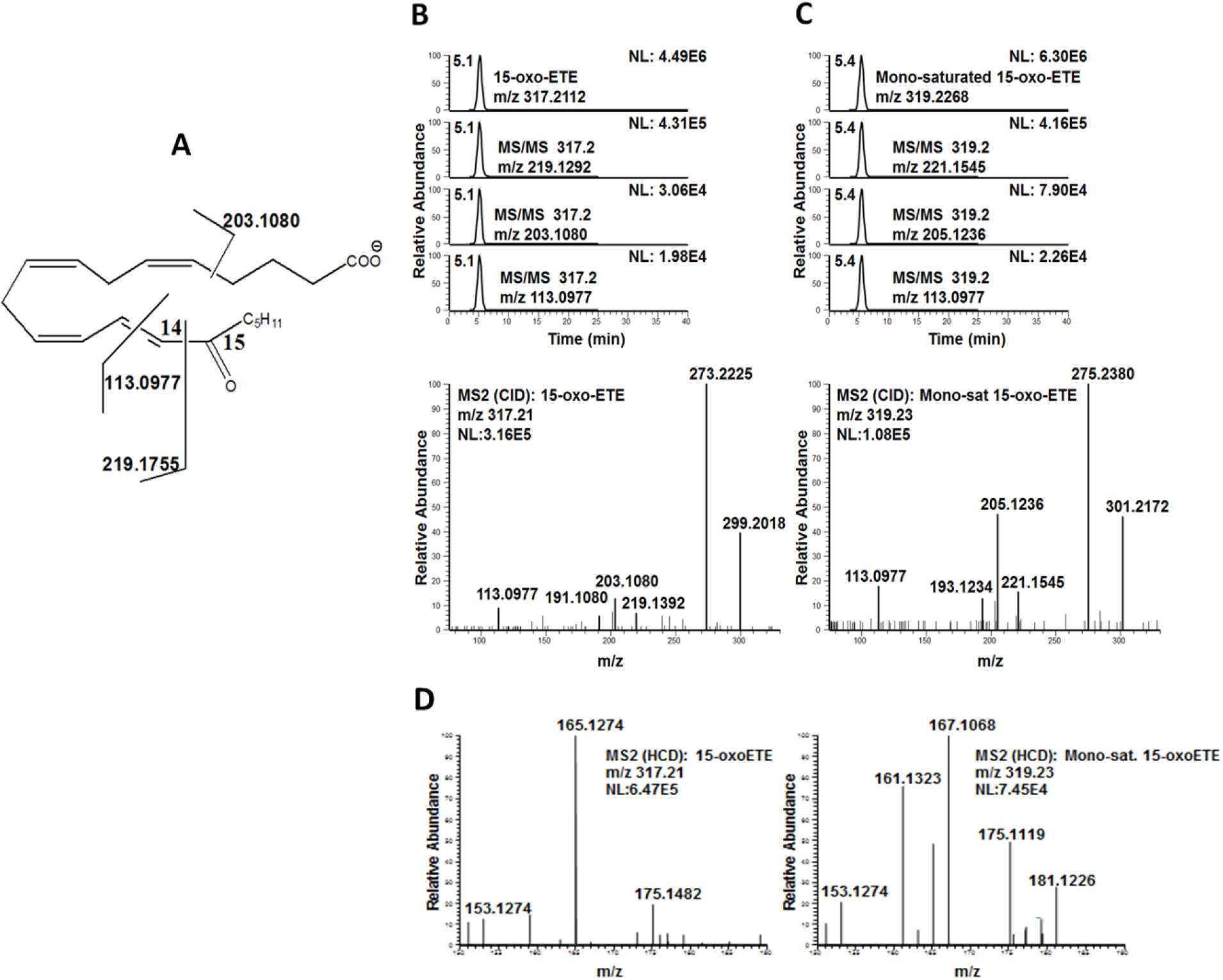
LC-MS/HRMS confirmation of the 15-oxoETE saturation product. (A) 15-oxoETE structure and diagnostic fragmentation. Extracted chromatograms and product ion spectrum in negative ion mode for (B) 15-oxoETE and (C) the mono-saturated 15-oxoETE metabolite. (D) Production ion spectra of 15-oxoETE and the mono-saturated 15-oxoETE metabolite between *m/z* 150-190 to highlight the fragmentation found at *m/z* 165.1274 for 15-oxoETE and *m/z* 167.1068 for the mono-saturated 15-oxoETE metabolite. The m/z of 167.1068 corresponds to fragmentation the chemical composition C_10_H_15_O_2_, a fragment adjacent to the C8-C9 double bond that would likely not be present if saturation occurred at this position.

### 3.4 Michael addition adducts reveal the position of double bond reduction

To further elucidate the points of saturation for 11- and 15-oxoETE, we examined the formation of GSH-adducts from treated HUVECs by LC-HRMS. GSH-adducts were observed for the parent and mono-saturated 11-oxoETE (Fig. 5A) and the parent and mono-saturated 15-oxoETE (Fig. 5B). No GSH-adduct was observed for the di-saturated 11-oxoETE metabolite, confirming C12-C13 double bond saturation and thus loss of the electrophilic α, β-unsaturated carbonyl (Fig. 5A). The two peaks for the parent 11-oxoETE-GSH adduct correspond to the 1,4- and the 1,6-Michael addition products. Supporting the identification of the initial saturation of 11-oxoETE at the C14-15 double bond, the mono-saturated product displayed only one peak, which would be the case if the 1,6-Michael product could no longer form due to C14-15 double bond saturation (14, 29).

**Figure 5:**
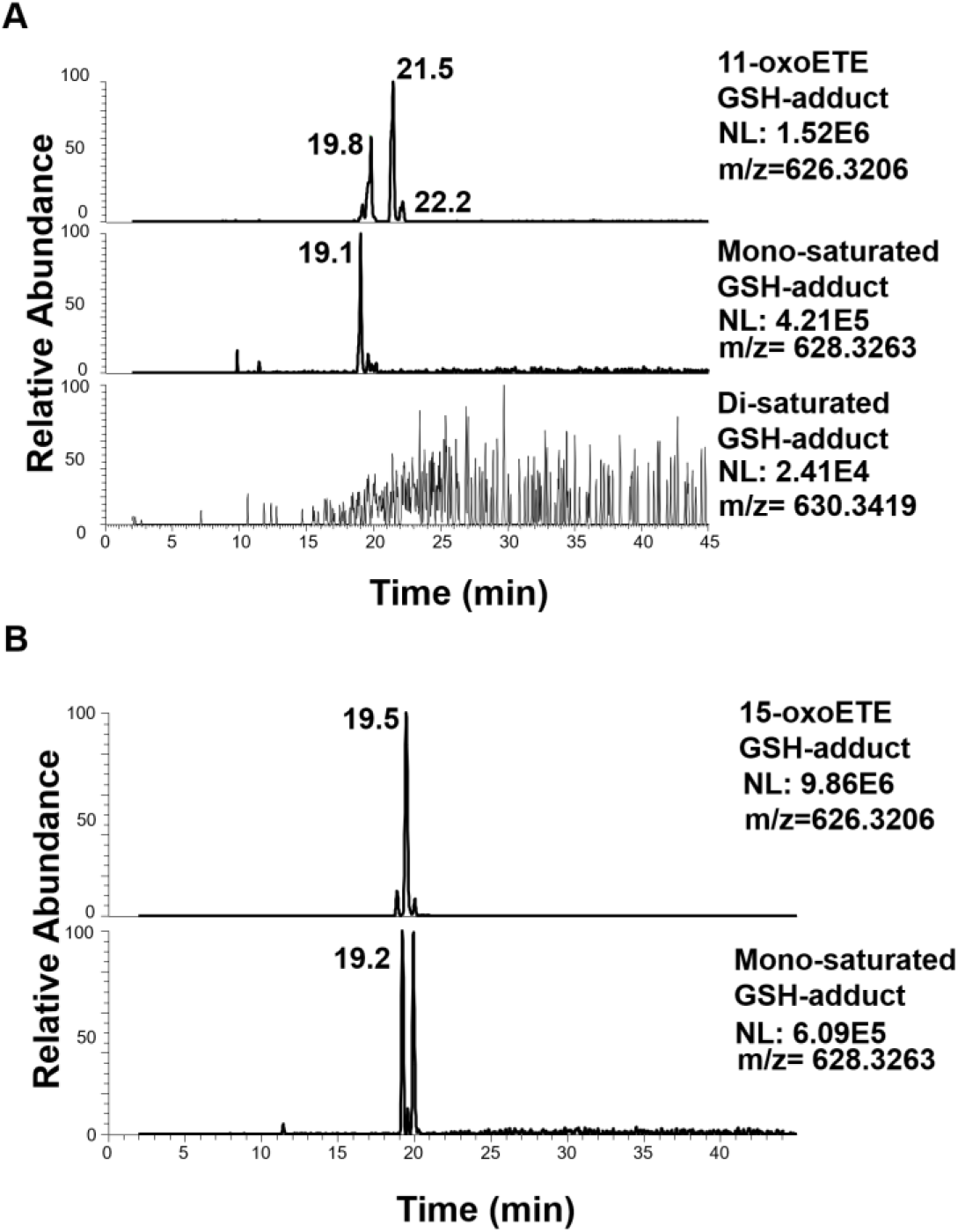
LC-HRMS chromatograms of GSH-adducts from parent and saturated products of 11-oxo and 15-oxoETEs. (A) 11-oxoETE, mono-saturated and di-saturated GSH conjugates (B) 15-oxoETE and its mono-saturated product.

### 3.5 Confirmation of 15-oxoETE metabolism in A549 cells

A549 cells were treated with 10 μM 15-oxoETE and samples were taken at time 0, 5 min, 30 min, 90 min, 3 hr and 6 hr for targeted LC-MS/MS analysis. The metabolism of 15-oxoETE to a mono-saturated metabolite was demonstrated to be the similar in A549 cells as the HUVECs (Fig. 6A). Formation of the mono-saturated 15-oxoETE metabolite occurred rapidly within the first 1.5 hr and steadily climbed until 3 hr where it plateaued Cell lysates were analyzed using high resolution MS on a Thermo Q Exactive coupled to a Vanquish UHPLC using the same gradient as the initial analysis to further characterize the mono-saturated species using parallel reaction monitoring. The fragmentation seen in A549 cells corresponds to that seen in HUVECs with the primary saturation taking place at the non-electrophilic double bond (Fig. 6B and C).

**Figure 6:**
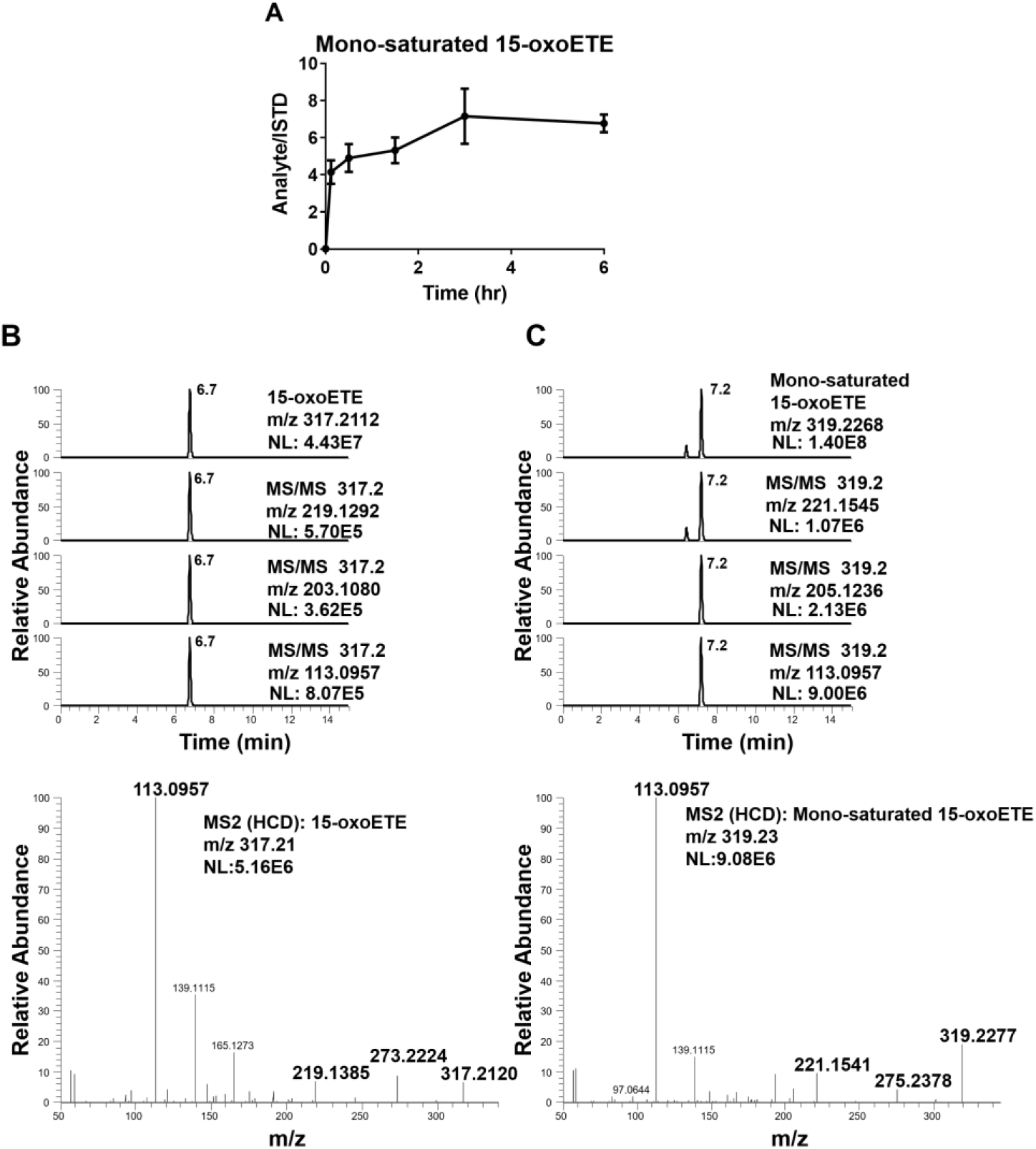
15-oxoETE metabolism in A549 Cells. (A) 15-oxoETE metabolism to the mono-saturated metabolite from 0 to 6 hr reported as SEM, n = 9 per condition. Extracted chromatograms of diagnostic product ions for (B) 15-oxoETE and (C) the mono-saturated 15-oxoETE metabolite.

### 3.6 The mono-saturated metabolite of the Ω-3 fatty acid 17-oxoDHA retains electrophilicity

A549 cells were exogenously administered 10 μM 17-oxoDHA, the oxo derivative of the Ω-3 fatty acid, docosahexaenoic acid. Treatment followed the same time course described above and formation of the mono-saturated metabolite peaked around 3 h in cell lysate (Fig. 6A). The mono-saturated product of 17-oxoDHA was characterized using LC-HRMS on the Q Exactive. The primary point of 17-oxoDHA saturation is more readily distinguished as the major fragment *m/z* 111.0801 (Fig. 7B) is shifted to 113.0957 (Fig. 7C). This *m/z* corresponds to fragmentation across the C7-C8 position indicating that saturation has occurred at the Ω-3 position and not at the α,β-unsaturated double bond. These results demonstrate that this metabolism is not unique to the arachidonic acid-derived oxoETE metabolites or to HUVEC cells.

**Figure 7:**
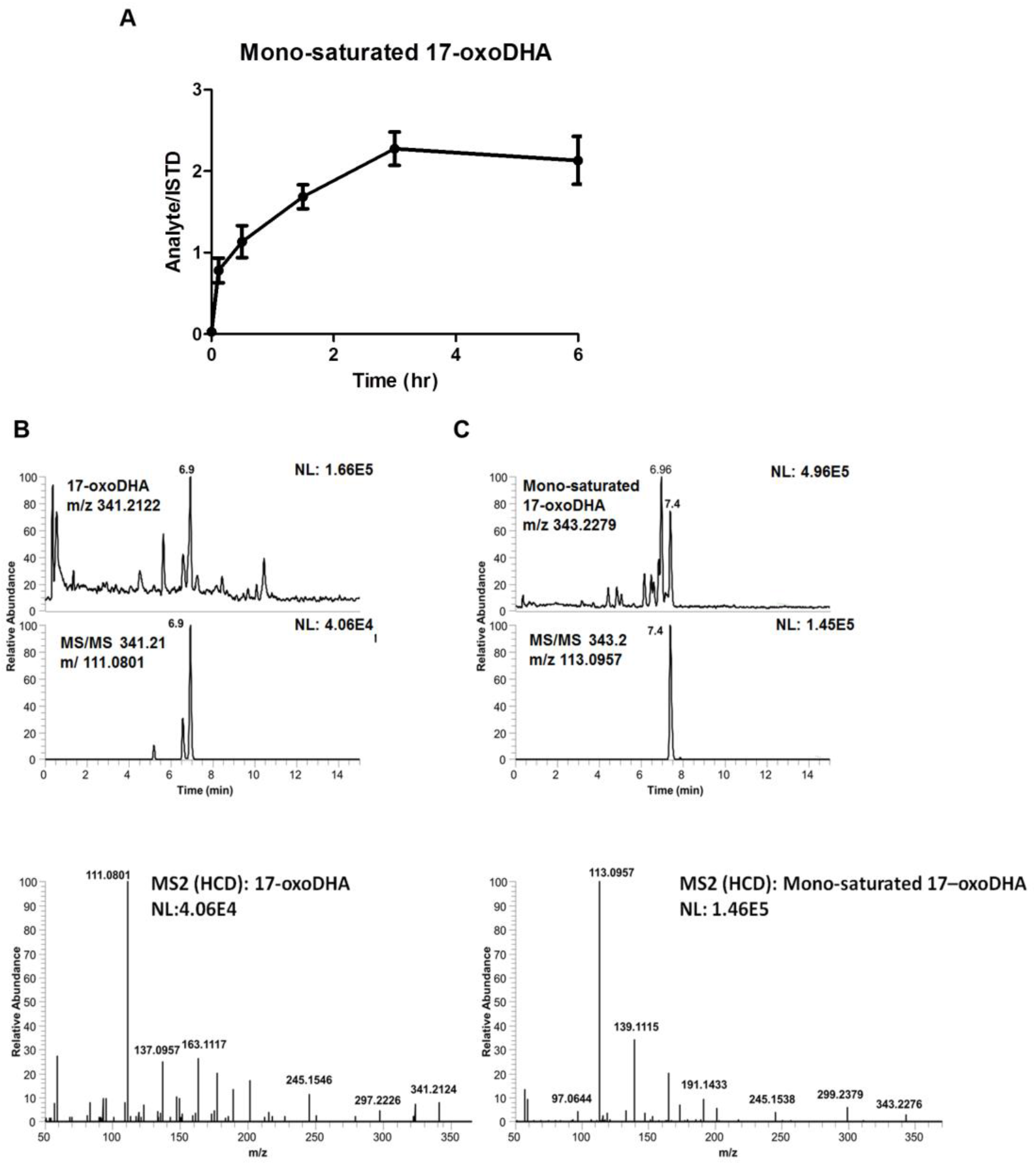
17-oxoDHA metabolism in A549 Cells. (A) 17-oxoDHA metabolism to the mono-saturated metabolite from 0 to 6 h reported as SEM, n = 9 per condition. Extracted chromatograms of diagnostic product ions and product ion spectra for (B) 17-oxoDHA and (C) the mono-saturated 17-oxoDHA metabolite.

### 3.7 Prostaglandin reductase saturates non-electrophilic double bonds

There are numerous dehydrogenases and reductases that may be responsible for the saturation of fatty acid double bonds (30). Prostaglandin reductase, PTGR1, is best known for the saturation of the α,β-double bond of 15-ketoPGE_2_, thus forming the non-electrophilic metabolite, 13,14-dihydro-15-ketoPGE_2_. Due to the similarity in structure between PGE_2_ and the oxoETEs it would seem plausible that PTGR1 would also reduce the α,β-unsaturated double bond of 15-oxoETE, 11-oxoETE and possibly even 17-oxoDHA. To test the potential of PTGR1 to reduce the double bonds in the oxoFAs, we performed an *in vitro* assay using recombinant purified PTGR1, reducing equivalents, and 11-oxo, 15-oxoETE, 17-oxoDHA, and 15-ketoPGE_2_ as substrates. Formation of the mono- and varying amounts of the di-saturated products derived from 11-oxo, 15-oxoETE, and 17-oxoDHA were observed with the combination of substrate, PTGR1 and NAD(P)H (Fig. 8). As expected, PTGR1 generated 13,14-dihydro-15-ketoPGE_2_ from 15-ketoPGE_2_ (Fig. 8 A and E). No artifactual saturation was observed without the enzyme present.

**Figure 8.**
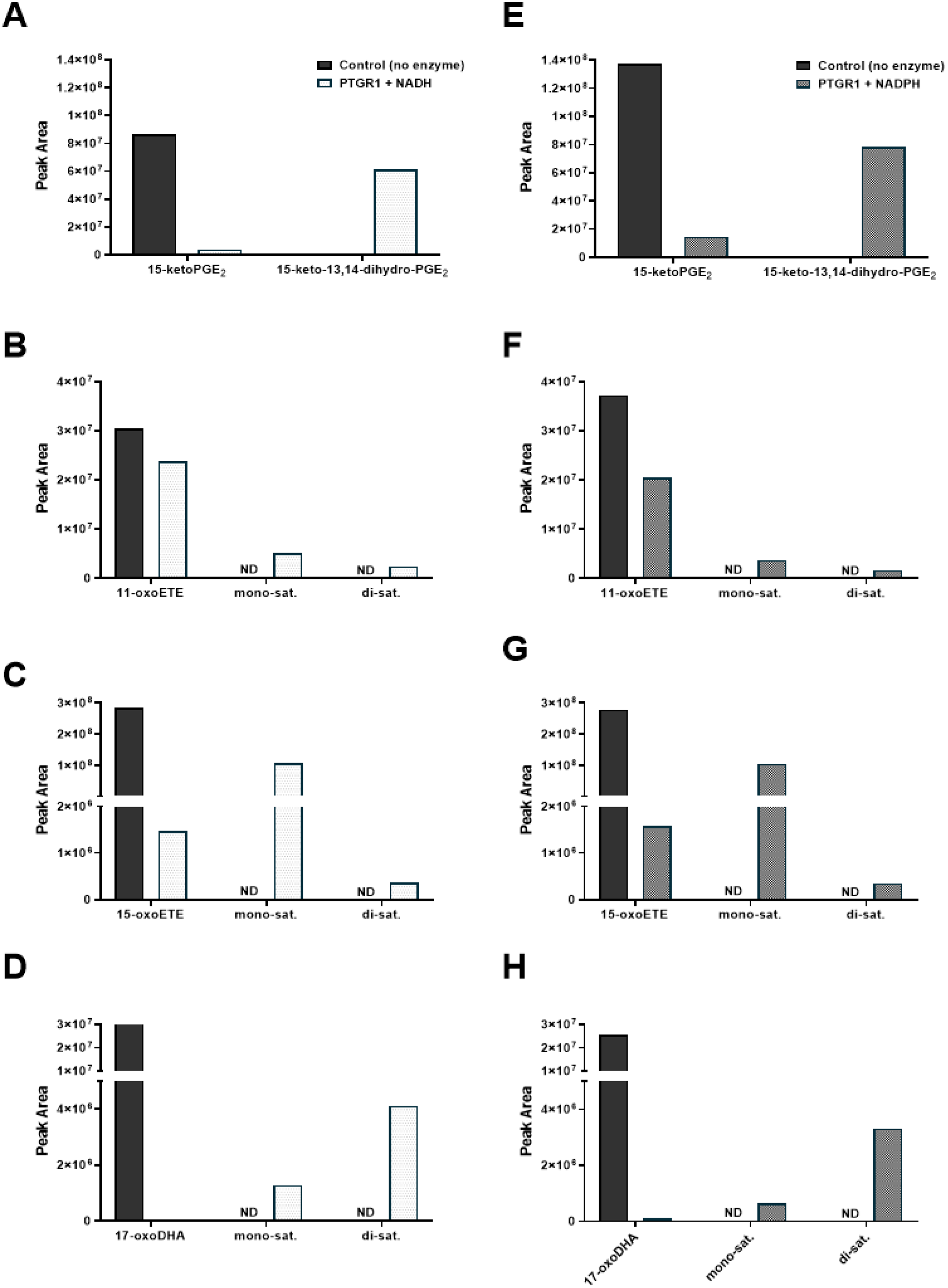
Purified recombinant PTGR1 catalyzes the conversion of oxoFAs to their saturated products. (A) 15-ketoPGE_2_ (B) 11-oxoETE (C) 15-oxoETE (D) 17-oxoDHA were converted to their saturated metabolites in the presence of PTGR1 and NADH (A-D) but not in reaction buffer without enzyme. Similar results were obtained with NADPH instead of NADH (E-H).

In order to investigate this role for PTGR1 in cells, we used lentiviral shRNA to knockdown PTGR1 protein expression via RNA interference (Table 1). A549 cells were transduced with lentivirus expressing control (Scr, scrambled shRNA) or PTGR1-specific shRNA. Western blot analysis indicates a ~50% knockdown (KD) of PTGR1 protein expression as compared to control and Scr (Fig. 9A). PTGR1-KD, Scr, and control A549 cells were treated with 10 μM 15-oxoETE and metabolites were analyzed using targeted analysis on the Sciex 6500 QTrap. No differences in 15-oxoETE metabolism were observed between A549 WT and Scr after 15-oxoETE treatment. However, PTGR1-KD cells displayed a significant decrease in the ratio of the mono-saturated metabolite/15-oxoETE at 3 and 6 h (Fig. 9B). Similarly, a significant decrease in the ratio of mono-saturated/17-oxoDHA was observed at 3and 6 h (Fig. 9C). 15-ketoPGE_2_ was used as the positive control and a significant decrease in 13,14-dihydro-15-ketoPGE_2_ was observed starting at 30 min and continued throughout the rest of the experiment (Fig. 9D). While PTGR1 does appear to play a role in 15-oxoETE and 17-oxoDHA metabolism, the decrease in the formation of the mono-saturated species in the PTGR1-KD cells is not as significant as 15-ketoPGE_2_ suggesting that other reductases are also capable of saturating double bond positions alternative to the electrophilic α, β double bond.

**Figure 9:**
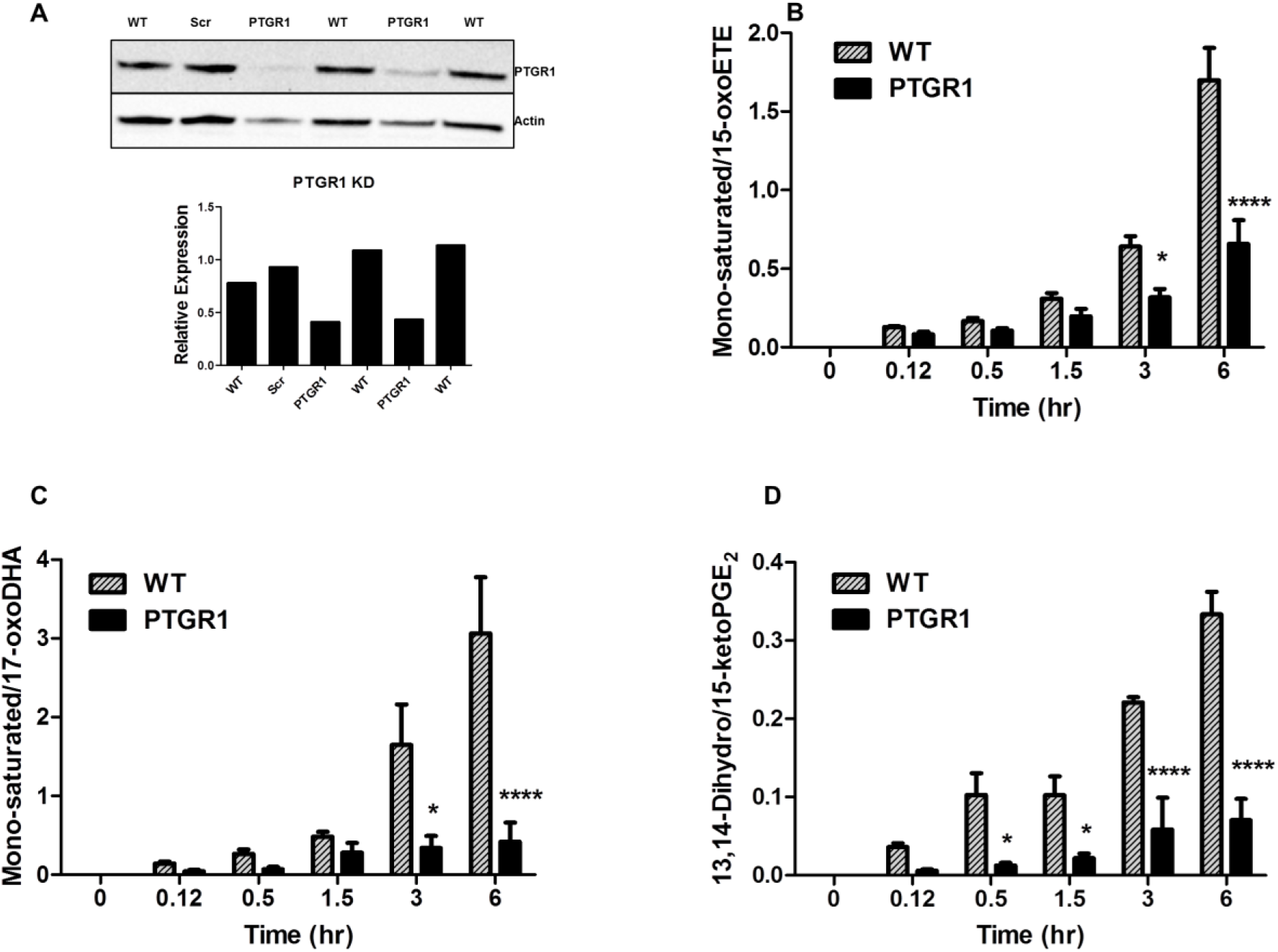
PTGR1 KD in A549 cells partially abolishes electrophile metabolism. (A) PTGR1 protein expression via western blot and image quantification Mono-saturated metabolite product/substrate ratio in A549 cell lysate for (B) 15-oxoETE, (C) 17-oxoDHA and the positive control (D) 15-ketoPGE_2_, n = 3-9 per condition, * p ≤ 0.05, **** p ≤ 0.0001.

Finally, we examined the formation of oxoFA metabolites using isogenic cell lines engineered for overexpression of PTGR1-FLAG (PTGR1-OE) as compared to empty vector controls. Western blots demonstrated a strong increase in PTGR1 expression (Fig 10A). Consistent with the analysis of the PTGR1 knockdown cells, PTGR1-OE decreased intracellular concentrations of the parent metabolite (Fig. 10 B, D, F), but increased the concentration of the mono-saturated products (Fig. 10 C E, G) following treatment with 10 μM of the parent oxoFAs. The overexpression did not significantly alter extracellular oxoFA metabolites (SFig. 1). Similar to the KD of PTGR1, these results indicate that PTGR1 is capable of saturating non-electrophilic double bonds.

**Figure 10.**
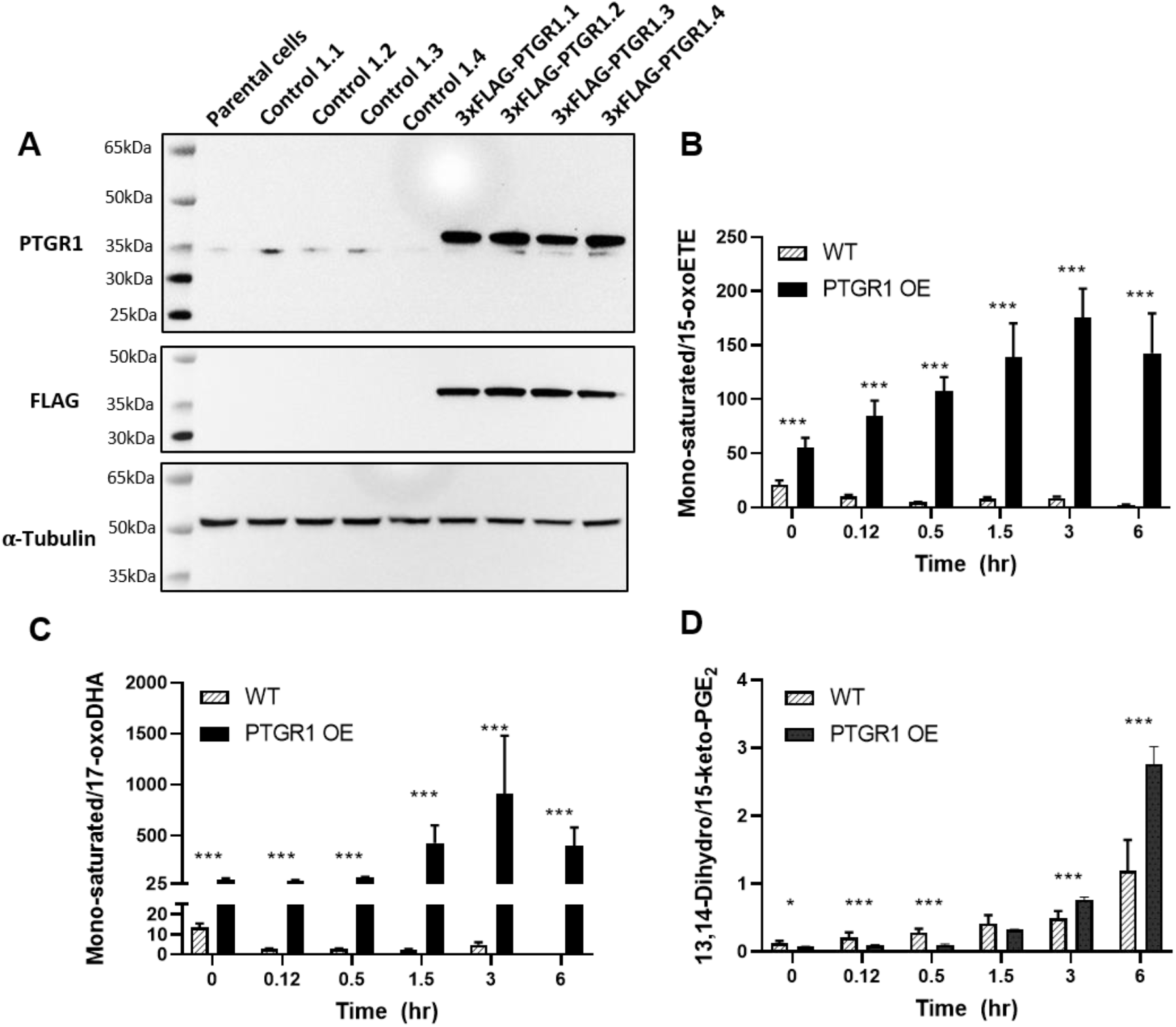
Overexpression of PTGR1 increases intracellular concentration of saturated oxoFAs. (A) PTGR1 protein expression (top) with FLAG tag (middle) and loading control (bottom). Mono-saturated metabolite product/substrate ratio in cell lysate for (B) 15-oxoETE, (C) 17-oxoDHA and the positive control (D) 15-ketoPGE_2_. n = 9 per condition, * p ≤ 0.05, *** p ≤ 0.001.

## 4. Discussion

The study of fatty acid metabolism has mostly focused on the formation of first generation bioactive lipids (i.e. HETEs, leukotrienes, PGs) and their signaling abilities, while downstream metabolism of these species remains poorly characterized. Existing literature on eicosanoid metabolism for COX/15PGDH products has focused on PGE_2_ and its 15PGDH-dependent electrophilic product 15-ketoPGE_2_, previously thought to be inactive; however, numerous studies including those focused on nitro-fatty acids have overwhelmingly demonstrated that electrophilic fatty acids are conferred with redox signaling capabilities (6, 8, 19, 31, 32). Further metabolism of 15-ketoPGE_2_ occurs via double bond reduction by PTGR1 to form the non-electrophilic product 13,14-dihydro-15-keto-PGE_2_. The electrophilic fatty acids 11- and 15-oxoETE are also products of 15PGDH and it would seem logical that reduction to a non-electrophilic metabolite would follow the same pathway as 15-ketoPGE_2_. However, this was not the case for either of these AA-derived bioactive lipids 11-oxoETE and 15-oxoETE, nor the DHA-derived electrophile, 17-oxoDHA (Scheme 1). In this work, we expand the knowledge of these pathways to include additional downstream products.

Our untargeted metabolomics approach revealed a distinct pattern of limited saturation for 11- and 15-oxoETE. The similar metabolism of [^13^C_20_]-15oxoETE, as well as time course studies support that these metabolites are derivatives of the parent oxoETEs. By examining the formation of GSH-derived Michael adducts we were able to discern the electrophilic status of the double bond proximal to the carbonyl in the α,β-unsaturated ketone moiety of the oxoETEs. The parent and predominant first saturation product of 11- and 15-oxoETE maintain electrophilicity, hence the α,β-unsaturated ketone moiety. However, the second saturation of 11-oxoETE no longer forms a GSH adduct, indicating the double bond at C12-C13 adjacent to the carbonyl has been reduced. The enzymatic basis of this transformation warrants investigation as the α,β-unsaturated ketone moiety has been associated with bioactivity in related molecules. These results were recapitulated in A549 cells and were also found to be relevant to the metabolism of the Ω-3 metabolite, 17-oxoDHA, which is also a bioactive lipid capable of activating the Nrf2 antioxidant response and inhibiting NF-κB-derived pro-inflammatory signaling (16).

To determine if PTGR1 plays a role in the metabolism of oxoETEs and oxoDHA species, we transduced A549 cells with lentivirus expressing control or PTGR1-specific shRNA and monitored the formation of saturated metabolites in cell lysate. Western blot demonstrated a 50% knockdown of PTGR1 protein expression in A549 cells and 15-ketoPGE_2_ was used as a positive control in the cell metabolism studies. PTGR1 KD significantly decreased formation of the mono-saturated metabolite when 15-oxoETE and 17-oxoDHA substrates were used, although not until 3 hr, whereas saturation of the positive control, 15-ketoPGE_2_ was significant as early at 30 min (Fig. 9). At the later time points, inhibition of saturation in PTGR1-KD cells was similar. 15-ketoPGE_2_ conversion to 13,14-dihydro-15-oxoPGE_2_ was decreased by ~4.4-fold at 3 hr, whereas formation of the mono-saturated metabolite decreased ~2-fold and ~4.9-fold for 15-oxoETE and 17-oxoDHA, respectively. This implies that other reductases have similar preferences for the double bond reduction of these fatty acid metabolites and may compensate for the loss of PTGR1 (Scheme 1).

Critical examination of oxo-fatty acid metabolism is warranted in light of the acceptance of electrophiles as FDA-approved therapeutics (dimethyl-fumarate) (33) and the extensive number of studies published on electrophilic oxo- and nitro-fatty acid signaling (8, 10, 34). Thus, we sought out to characterize the primary metabolic products of oxo-fatty acids in order to better understand their normal and pathological functions. Metabolic pathway characterization will also help to facilitate identification of the enzyme(s) responsible for the saturation of the electrophilic double bond. A variety of distinct fatty acid metabolizing enzymes exist with varying specificity for substrate and reaction (7) and there are numerous short and medium chain dehydrogenases/reductases that are still classified as orphans (35, 36). Elucidation of the pathways of metabolism will also allow for the design of therapeutic interventions to modulate or mimic the effects of these molecules, and aid in the discovery of inhibitors or the design of stable derivatives. As an additional benefit, identification of these novel metabolites may contribute to the ongoing efforts in untargeted metabolomics to build high resolution databases as well as an increased appreciation of specificity in lipidomics experiments.

## abbreviations

15d-PGJ_2_: 15-deoxy-Δ^12,14^-PGJ_2_
15-PGDH: 15-prostaglandin dehydrogenase
AA: arachidonic acid
BME: β-mercaptoethanol
CID: collision induced dissociation
COX: Cyclooxygenase
GSH: glutathione
HPETE: hydroperoxyeicosatetraenoic acid
HUVECs: human umbilical vein endothelial cells
LO: lipoxygenase
ME: methyl ester
oxoETE: oxoeicosatetraenoic acid
PGE_2_: prostaglandin E_2_
EP: prostaglandin E receptor
PTGR1: prostaglandin reductase 1
SPE: solid phase extraction
XIC: extracted ion chromatogram

## Acknowledgements

This work was supported by the National Institutes of Health P30ES013508 (IAB), K22ES26235 (NWS), UL1TR000005 and R21AI12207 (Wendell). Research in the Sobol lab is funded by grants from the National Institutes of Health (NIH) [CA148629, ES029518, ES028949, CA238061, AG069740 and ES032522] and from the National Science Foundation (NSF) [NSF-1841811]. Support is also provided from the Abraham A. Mitchell Distinguished Investigator Fund and from the Mitchell Cancer Institute Molecular & Metabolic Oncology Program Develop fund (to RWS).

## Disclosures

RWS is a scientific consultant for Canal House Biosciences, LLC.

## Supplemental Figures

**Supplementary Figure 1.**
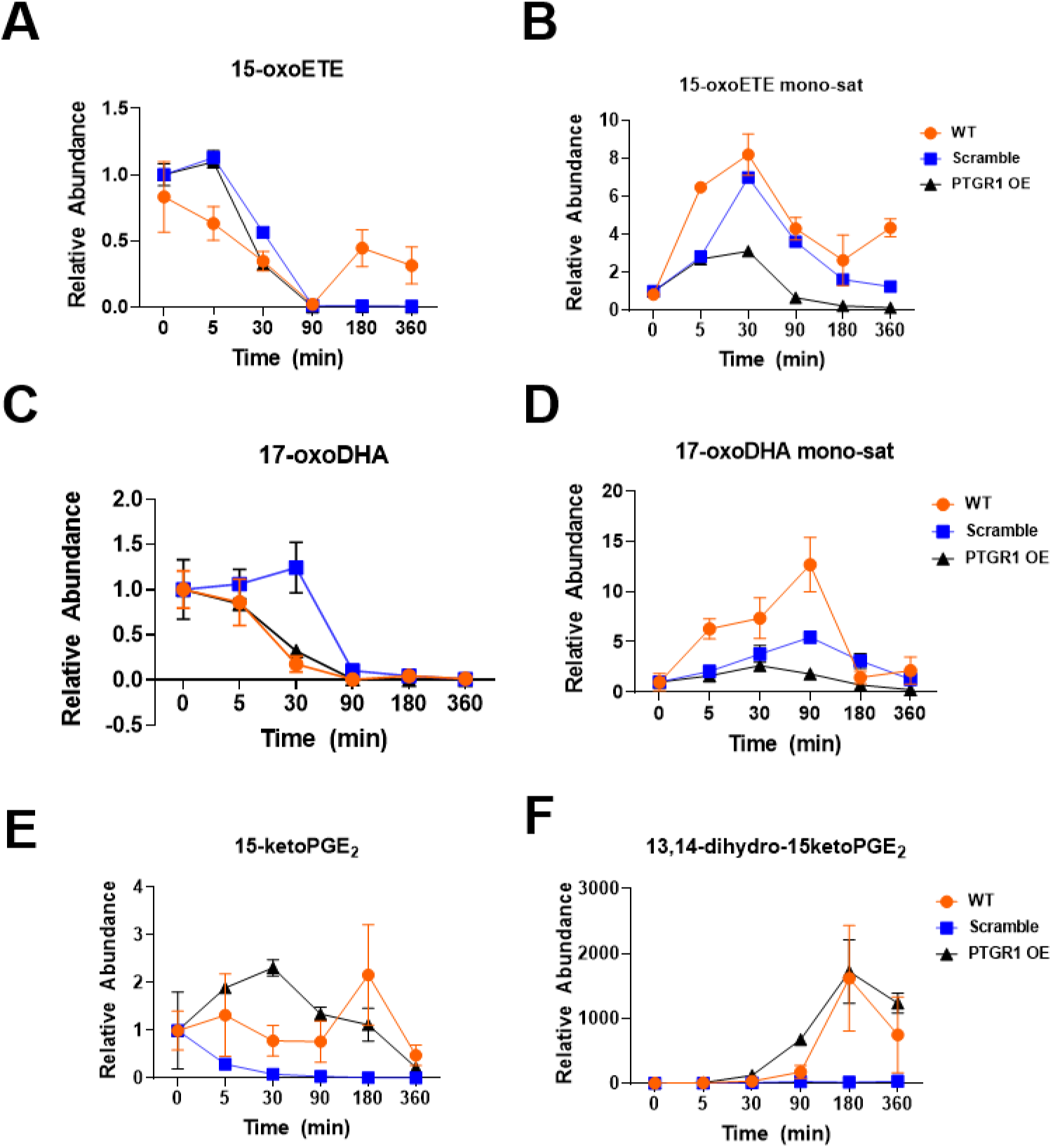
Overexpression of PTGR1 did not increase the extracellular concentration of saturated oxoFAs. PTGR1 OE in A549 cells does not result in an increase in the export of PUFA-derived lipid mediators, graphed as SEM.

**Scheme 1:**
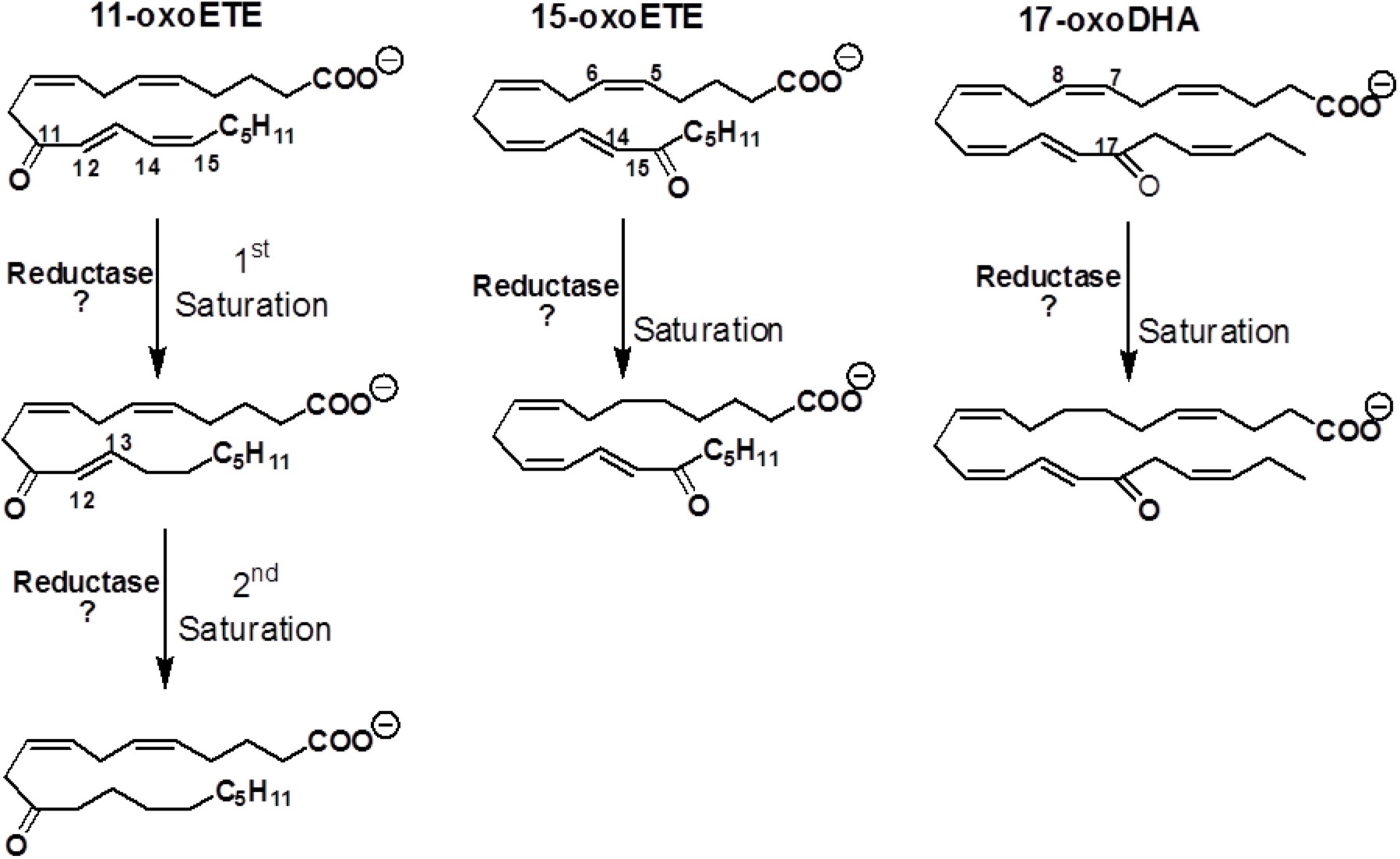
Structures of 11-oxoETE, 15-oxoETE, 17-oxoDHA and proposed saturation products. 11-oxoETE undergoes two rounds of saturation to form mono- and di-saturated metabolites whereas only mono-saturated metabolites were found for 15-oxoETE and 17-oxoDHA. Proposed putative points of saturation based off of the MS^2^ spectra are also shown.

## Notes

### Competing Interest Statement

SGW declares interest in Complexa Inc. RWS is a scientific consultant for Trevigen, Inc.

### Summary of Updates

Additional experiments (overexpression and purified enzyme experiments).

